# Rapid intracellular acidification is a novel plant defense response countered by the brown planthopper

**DOI:** 10.1101/2023.05.24.542203

**Authors:** Yanjuan Jiang, Xiao-Ya Zhang, Shaoqin Li, Yu-Cheng Xie, Xu-Mei Luo, Yongping Yang, Zhengyan Pu, Li Zhang, Jia-Bao Lu, Hai-Jian Huang, Chuan-Xi Zhang, Sheng Yang He

## Abstract

The brown planthopper (BPH) is the most destructive insect pest in rice. Through a stylet, BPH secrets a plethora of salivary proteins into rice phloem cells as a crucial step of infestation. However, how various salivary proteins function in rice cells to promote insect infestation is poorly understood. Among them, one of the salivary proteins is predicted to be a carbonic anhydrase (NlCA). The survival rate of the *NlCA-*RNA interference (RNAi) BPH insects was extremely low on rice, indicating a vital role of this salivary protein in BPH infestation. We generated *NICA* transgenic rice plants and found that *NlCA* expressed in rice plants could restore the ability of *NlCA-*RNAi BPH to survive on rice. Next, we produced rice plants expressing the ratiometric pH sensor pHusion and found that *NlCA-*RNAi BPH induced rapid intracellular acidification of rice cells during feeding. Further analysis revealed that both *NlCA-*RNAi BPH feeding and artificial of lowering intracellular pH activated plant defense responses, and that NICA-mediated intracellular pH stabilization is linked to diminished defense responses, including reduced callose deposition at the phloem sieve plates and suppressed defense gene expression. Given the importance of pH homeostasis across the kingdoms of life, discovery of NICA-mediated intracellular pH modulation uncovered a new dimension in the interaction between plants and piecing/sucking insect pests. The crucial role of NICA for BPH infestation of rice suggests that NICA is a promising target for chemical or trans-kingdom RNAi-based inactivation for BPH control strategies in plants.

**Highlights:**

- Rapid intracellular acidification is a novel plant defense response.
- Brown planthopper delivers into rice tissues a major virulence protein, carbonic anhydrase (*Nl*CA), critical for survival.
- NlCA counters intracellular acidification to suppress plant defense.
- Results implicate existence of pH-responsive regulators of defense responses in the rice-brown planthopper interaction.

**In brief:** The brown planthopper (BPH) is the most devasting insect pest in rice. Jiang *et al*. discovered that BPH secretes a salivary carbonic anhydrase (*NI*CA) to regulate the intracellular pH of the rice cell to facilitate its feeding and survival on rice plants and that NICA-counters host intracellular pH acidification to diminished defense responses. These findings uncovered that intracellular pH homeostasis is a previously uncharacterized battleground in plant-insect interactions, but also open a door to future discovery of pH-responsive intracellular regulators of defense responses, which could add a new research dimension in the study of plant-biotic interactions.

## INTRODUCTION

The brown planthopper (BPH; *Nilaparvata lugens* Stål, Hemiptera, Delphacidae) is a monophagous insect pest of rice (*Oryza sativa* L.) found in all rice-growing Asian countries. The BPH sucks rice phloem sap via its stylet, causing leaf yellowing and wilting, stunted plant growth, reduced photosynthesis and ultimately death of rice plant^1^. During severe BPH outbreaks, tens of characterized by large-scale wilting, yellowing and lethal drying of rice plants^2^. Besides direct damages, the BPH may also indirectly damage rice plants by oviposition and transmitting viral disease agents^3,4^.

Application of chemical insecticides has been a main strategy for controlling BPH. Although it has the advantages of having rapid effects on killing insects and low costs, use of insecticides leads to environmental pollution and resistance of BPH to pesticides. In recent decades, breeding rice resistant varieties to control BPH has attracted increasing attention. To date, more than 30 resistance genes have been found in the rice genome^5^. However, BPH often quickly evolves new biological types that evade rice resistant genes. Therefore, additional BPH controlling methods need to be developed to complement the current control measures toward long-term solutions of achieving durable BPH resistance in rice. One method could be based on disruption of key steps of BPH’s natural infestation process. However, development of such methods will require a comprehensive understanding of the basic biology of the BPH-rice interaction.

A critical step in BPH infestation of rice is secreting bioactive substances into the plant tissues through the stylet^6–8^. Specifically, during feeding, BPH secretes both colloidal and watery saliva^9, 10^. The main function of the colloidal saliva is to form a saliva sheath around the piercing-sucking mouthparts, stabilizing the overall feeding apparatus. The composition and function of watery saliva is more complex, and it contains salivary proteins that are believed, in most cases, to regulate various pathways in plant cells to enhance BPH feeding and survival in rice plants. For example, NlEG1, a salivary endo-b-1,4-glucanase, degrades plant celluloses to help the BPH’s stylet reach to the phloem^11^. NlSEF1, an EF-hand Ca^2+^-binding protein, interferes with calcium signaling and H_2_O_2_ production during BPH feeding^12^. Salivary protein 7 is required for normal feeding behavior and for countering accumulation of a defense compound, tricin^13^. On the other hand, mucin-like protein (NlMLP) triggers defense responses in rice cells, including cell death, callose deposition and up-regulation of pathogen-responsive genes^14,15^.

Carbonic anhydrases (CAs) (EC 4.2.1.1) are zinc metalloenzymes that function as catalysts in the bidirectional conversion of CO_2_ and water into bicarbonate and protons^16^. There are at least five distinct CA families (α-, β-, γ-, δ-, and ε-CAs) and three of them (α-, β-, and γ-CAs) are ubiquitously distributed among animal, plant, and bacterial species. The widespread distribution and adequate abundance of these CA families underline their evolutionary importance throughout the kingdoms of life. CAs participate in a wide range of biological processes, such as pH regulation, CO_2_ homeostasis, stomatal aperture, and plant defense^17–21^. NICA belongs to the α-CA subfamily. Our previous study showed that *NlCA* expressed in BPH salivary glands^6^. Surprisingly, however, RNA interference (RNAi) of the *NlCA* transcript in BPH insects affects neither pH maintenance within the salivary gland, watery saliva or gut, nor insect feeding behavior or honeydew excretion, but greatly reduced survival of BPH on rice plants, suggesting a critical function *in planta* via an unknown mechanism^6^.

Here, we report that *NlCA-*RNAi BPH feeding results in rapid intracellular acidification of rice cells. We found that NlCA is secreted into the rice tissues and functions as an effector that stabilizes host cell intracellular pH, accompanied by suppression of defense responses, during BPH feeding. Thus, we have uncovered intracellular pH homeostasis is a previously uncharacterized battleground in plant-insect interactions.

## Results

### NlCA is detected in rice sheath tissues during BPH feeding

NlCA was previously found to be highly expressed in salivary glands and present in the watery saliva of BPH fed on artificial diet^6^. We conducted a more detailed characterization of *NICA* expression in this study. RNA *in situ* hybridization showed that the expression level of *NlCA* was detectable throughout the principal glands (PGs) and accessory glands (AGs), but not in A-follicle of the principal gland (APG) (Figure 1A), which further raised the possibility that NICA may be one of the “effector proteins” secreted into the rice tissue during BPH feeding on rice plants. To test this possibility, we compared protein profiles in the leaf sheaths of Nipponbare rice plants before and after BPH feeding using liquid chromatograph-mass spectrometer (LC-MS). We found 8 NICA-specific peptides in BPH-fed leaf sheath tissue (Figure 1B), confirming that NlCA is secreted into the host tissues during BPH feeding. As BPH is a phloem-feeding insect, its effector proteins, such as NlCA, presumably act in the phloem. We therefore collected the phloem exudate from Nipponbare rice leaf sheaths with or without BPH feeding for LC-MS analysis (Figure 1C). Six NlCA-specific peptides were detected in the phloem exudate of Nipponbare plants fed by BPH (Figure 1B), demonstrating that NlCA is delivered into phloem by BPH.

**Figure 1.**
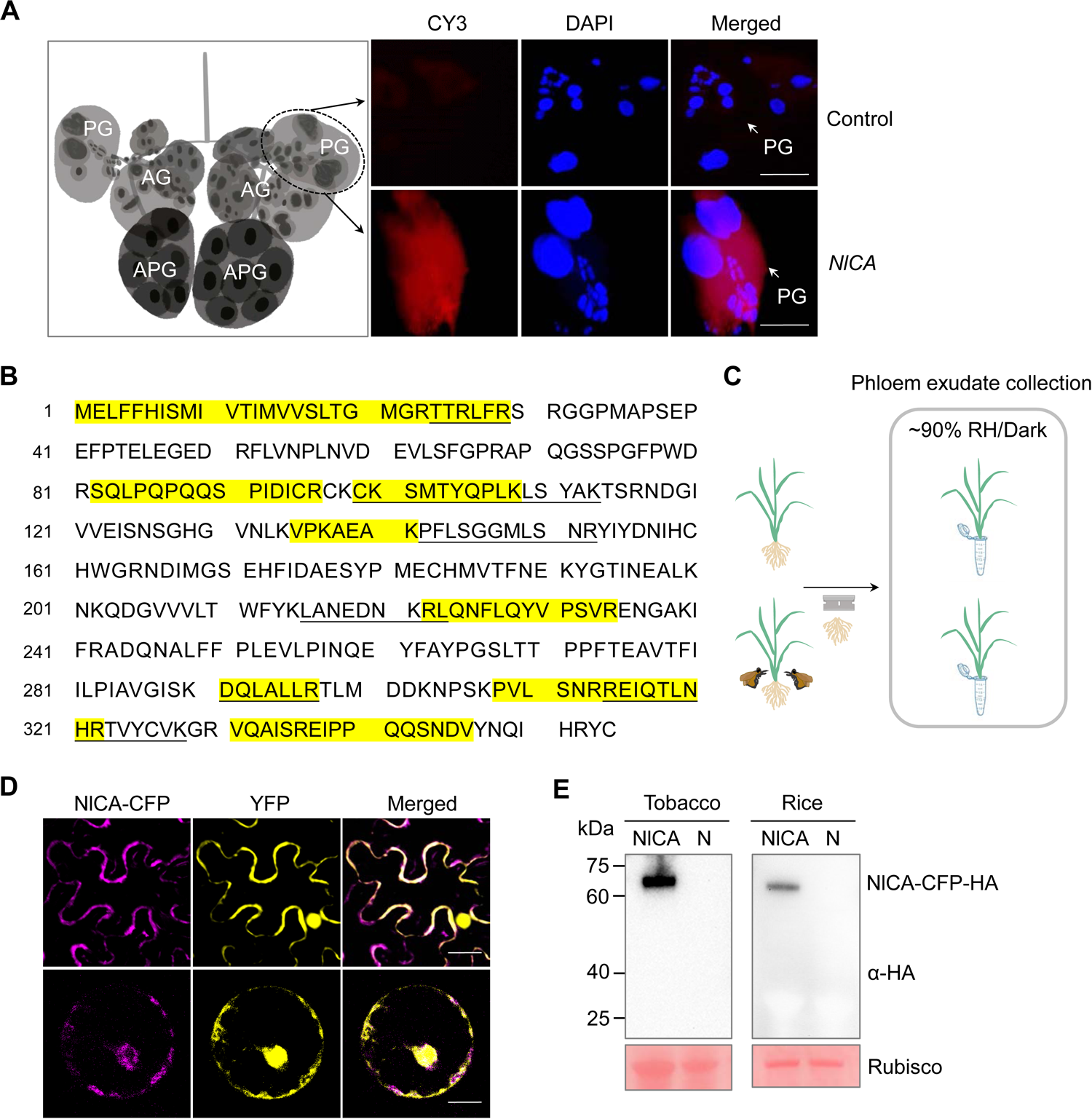
Initial characterization of *Nilaparvata lugens* carbonic anhydrase (NlCA). (A) *in situ* RNA hybridisation of salivary glands in 5-instar BPHs. Left, a schematic diagram of BPH salivary glands, including the principal glands (PG), the accessory glands (AG) and the A-follicle of the principle gland (APG). Right, *NlCA* expression was detectable in the PG (red) using antisense *NICA* sequence as a probe. Sense *NICA* probe was used as a negative control. Nuclei are stained blue by 4’,6-diamidino-2-phenylindole (DAPI). Scale bar = 50 μm. (B) The amino acid sequence of NlCA. The highlighted amino acid residues indicate the peptides detected in BPH-infested rice sheath tissue by LC-MS analysis. The underlined amino acid residues indicate the peptides detected in phloem exudate of BPH-infested rice by LC-MS analysis. (C) A schematic diagram of the phloem exudate collection. (D) NlCA is colocalized with the YFP signals in *N. benthamiana* (upper row; Scale bar = 10 μm) and rice cells (lower row; Scale bar = 5 μm). YFP and NlCA-CFP fusion proteins were co-expressed in *N. benthamiana* leaf cells for 48 h using the *Agrobacterium*-mediated transient expression method. YFP and NlCA-CFP fusion proteins were co-expressed in rice protoplasts 16 h after the corresponding DNA constructs were introduced into rice protoplasts via polyethylene glycol-mediated transformation. Experiments were repeated three times with similar trends. (E) NlCA-CFP-HA fusion protein levels are detected with anti-HA (Zenbio, 301113) in *N. benthamiana* leaves and rice protoplasts. Protein samples were extracted from NlCA-CFP-HA expressing *N. benthamiana* and rice protoplasts. Proteins from mock *N. benthamiana* leaves and rice protoplasts were as negative controls (N). Ponceau S staining of Rubisco shows protein loading control. See also Table S1.

### Transgenic expression of *NlCA* in rice rescues the ability of *NlCA*-silenced BPH to feed and survive

To further clarify the site of function (i.e., in insect vs. in plant) of NlCA in the BPH-rice interaction, we produced transgenic Nipponbare plants expressing *NICA* (see Methods). A total of 26 lines were produced and 6 lines were found to robustly express the *NICA* transcript (Supplemental Figure 1A). *NlCA*-expressing plants exhibited no noticeable changes in appearance compared to Nipponbare plants (Supplemental Figure 1B–1D). *NlCA*-expressing plant lines were propagated to T3 generation, and three lines were subjected to further characterization, including BPH feeding. For BPH feeding assay, double-stranded RNA (dsRNA) of *NlCA* (ds*NlCA*) or the control green fluorescent protein gene (ds*GFP*) was injected into 3^rd^ instar BPH nymphs to initiate RNAi of the *NICA* transcript^22^. Quantitative real-time PCR analysis confirmed that the transcript levels of the *NlCA* gene were reduced by 99% and 97%, respectively, in tested individuals when compared with non-RNAi control and ds*GFP*-treated insects (Figure 2A). There were no significant differences in the survival rates between ds*GFP* and ds*NlCA* BPH when fed on the artificial diet, indicating that silencing *NlCA* expression has no obvious impact on the basic physiology of BPH (Figure 2B). However, we found that the survival rate of ds*NlCA* BPH was sharply decreased, starting at day 9 post-infestation, to ∼40% at day 14, whereas the control ds*GFP* BPH survived normally on wild-type Nipponbare plants (Figure 2C). This result is consistent with previous results conducted in the *japonica* cultivar Xiushui134 rice plants^6^, suggesting that the requirement of NICA for BPH survival is not specific to a specific rice genotype. Strikingly, *NICA* transgenic plants almost fully restored the survival of *NlCA*-silenced BPH insects (Figure 2C; Supplemental Figure 1E–1F), demonstrating that *Nl*CA expressed in the host tissue (Figure 2D and 2E) can complement the infestation defect of *NlCA*-silenced BPH insects.

**Figure 2.**
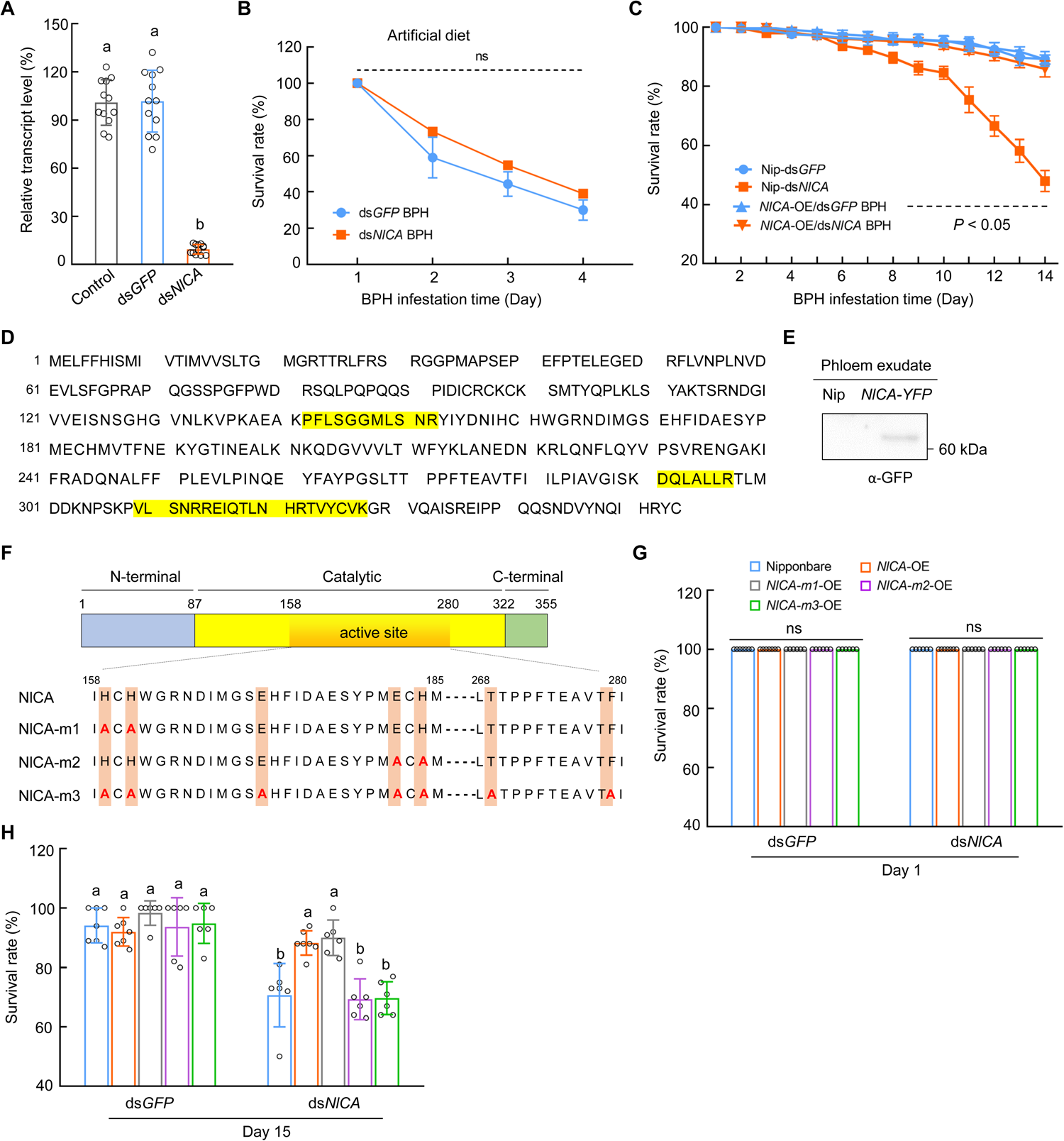
Effects of *NlCA* RNAi on BPH survival on rice cultivar Nipponbare. (A) The *NICA* transcript levels in BPH at 3 dpi post-injection of ds*GFP* or ds*NICA* were determined by qRT-PCR (displayed as % of the *NICA* transcript abundance in control BPH). Values are displayed as mean ± SEM of 4 experimental replicates (Two-way ANOVA; 3 biological replicates in each experiment and 30 individual insects pooled for each biological replicate). (B) The daily survival rates of ds*NlCA*-injected BPH insects feeding on artificial diet. Values are displayed as mean ± SEM of 3 biological replicates (one biological replicate includes 50 individual BPH adults fed on artificial diet in a lucifugal plastic bottle). (C) The daily survival rates of ds*NlCA*-injected BPH insects feeding on Nipponbare and *NlCA*-expressing line 1. Values are displayed as mean ± SEM of 8 biological replicates (Two-way ANOVA; 20 individual 2nd BPH nymphs per rice plant for each biological replicate). The black dashed line indicates significant differences (*P* < 0.05) in survival rates start day 9 between ds*NICA* BPH on Nipponbare (Nip) and other treatments. (D) The amino acid sequence of NlCA. The highlighted amino acid residues indicate the peptides detected in phloem exudate of *NlCA*-OE transgenic rice plants by LC-MS analysis. (E) NlCA:YFP protein is detectable in the phloem exudate of *NlCA*-OE transgenic plants by anti-GFP antibody (Abmart, 20004). (F) Schematic display of the NlCA protein with the N-terminus displayed in bule, the C-terminus in green and the central catalytic domain in yellow/orange. The amino acid sequence (158–280 aa) of the putative active site is showed. Amino acid substitutions in NlCA-m1, NlCA-m2 and NlCA-m3 are indicated in red. (G and H) The survival rates of 20 ds*GFP* and ds*NlCA* BPH insects in Nipponbare and *NICA* transgenic plants after 1-day (G) and 15-days (H) post-infestation. Values are displayed as mean ± SEM (n ≥ 6 biological replicates; 20 individual 2^nd^ BPH nymphs per rice plant for each biological replicate). ns indicates no significant difference between treatments (Two-way ANOVA). Different letters indicate statistically significant differences analyzed by two-way ANOVA (Tukey test, *P* < 0.05). Experiments were repeated three times with similar trends. See also Figure S1.

### The conserved catalytic site amino acids of *Nl*CA are critical to its function in rice

NlCA contains several conserved amino acid residues predicted to be at the catalytic site of carbonic anhydrases (Figure 2F). We asked if some of these conserved active site residues are critical to the function of *Nl*CA in the rice-BPH interaction. Accordingly, we expressed three different *NlCA* mutants in Nipponbare plants (Supplemental Figure 1A and 1G). “*NlCA-m1*” plants contain a *NICA* transgene in which His^159^ and His^161^ were mutated to Ala; “*NlCA-m2*” plants contain a *NICA* transgene in which Glu^182^ and His^184^ were replaced by Ala; and “*NlCA-m3*” plants carry a *NICA* transgene in which all seven conserved active site residues, His^159^, His^161^, Glu^171^, Glu^182^, His^184^, Thr^269^ and Phe^279^, were changed to Ala. Like *NICA* transgenic plants, transgenic plants expressing these *NlCA*-mutants exhibited no noticeable changes in appearance (Supplemental Figure 1B-1D). We conducted BPH survival assay and found that, in contrast to plants expressing wild-type *NICA*, the *NlCA-m2* and *NlCA-m3* plants could not fully rescue the infestation defect of *NlCA*-silenced BPH insects. The *NlCA-m1* plants, on the other hand, could recover the infestation defect of ds*NICA* BPH (Figure 2G and 2H; Supplemental Figure 1H). Thus, the conservative residues Glu182 and His184 at the predicted catalytic site are indispensable for the function of *Nl*CA inside the plant cell.

After showing the requirement of catalytic site residues for the function of NICA in rice, we next investigated the subcellular localization of NICA in both *Nicotiana benthamiana* leaf cells and rice leaf protoplasts transiently expressing *NlCA:CFP*. In both cases, NlCA:CFP showed a nonuniform distribution of CFP signal in the cell. Further colocalization studies with YFP signal revealed that NICA was predominantly localized in the cytoplasm (Figure 1D). There was some nuclear localization of NlCA-CFP in rice protoplasts. Intact NlCA:CFP proteins were detected in these localization experiments (Figure 1E).

### NlCA counters rapid intracellular acidification during BPH feeding of rice

BPH feeding on rice plants can be divided into two phases based on the electropenetrography (EPG) waveforms^23^. The first phase involves BPH’s stylet penetrates the plant through the cell walls and cell membranes of various epidermal and mesophyll cells until the stylet reaches the phloem sieve cells inside the vascular system. In Nipponbare, the time to reach the phloem sieve cells can be around 1.2 h before sustained phloem sap ingestion occurs at 3.8 h^23^. Our finding of the site of NICA action in the plant prompted us to test the hypothesis that NICA might modulate rice cell pH changes during BPH feeding. To directly test this hypothesis, we constructed a rice transgenic line expressed a cytoplasmic ratiometric pH sensor driven by the 35S promoter, cyto-pHusion. Cyto-pHusion consists of the tandem concatenation of enhanced green fluorescent protein (EGFP) and a monomeric red fluorescent protein 1 (mRFP1)^24^. EGFP is highly sensitive to pH variation with the brightest EGFP signal emitted at a pH of 7–8. EGFP fluorescence is gradually quenched at lower pH values and totally quenched at pH values <5. In contrast, mRFP1 is insensitive to pH changes in the physiologically relevant range and serves as an internal reference. We placed 5^th^ instar BPH nymphs on the leaf sheath of cyto-pHusion-expressing rice plants and recorded GFP and RFP signals of the feeding sites under microscopic observation at different time points with a 12 h test period. BPH can move around from one feeding site to another during the 12-h feeding experiments. The average feeding time at a feeding site is around 8 h. Therefore, the first 8 h of cellular responses best capture mostly synchronized rice-BPH interactions at a typical feeding cycle. As shown in the Figure 3A–D, ds*NlCA* BPH feeding induced a significant pH decrease in the phloem sieve elements/companion cells at 4 h and 8 h (Figure 3C and D). The decreased ratio of EGFP: mRFP was caused by the quenched EGFP signals after ds*NlCA* BPH feeding, while the internal control, mRFP, kept at a stable level (Supplemental Figure 2A–2H). In contrast, ds*GFP* BPH feeding did not elicit an obvious intracellular pH change in the phloem sieve elements/companion cells during this period. These results uncovered a previously uncharacterized plant cellular response, rapid intracellular acidification, during the early stage of ds*NICA* BPH feeding and demonstrated that BPH has evolved a critical virulence effector, NlCA, to counter this novel plant cellular response. Interestingly, we noted that the control ds*GFP* wild-type BPH feeding induced slight intracellular acidification at 12 h (Figure 3E). At this late time point of feeding, most BPHs would have left their initial feeding sites and have relocated to new feeding sites. In our analysis, we could not distinguish between old and new feeding sites. It is therefore likely that active BPH feeding (i.e., active NlCA injection) is needed to maintain intracellular pH homeostasis at all feeding sites.

**Figure 3.**
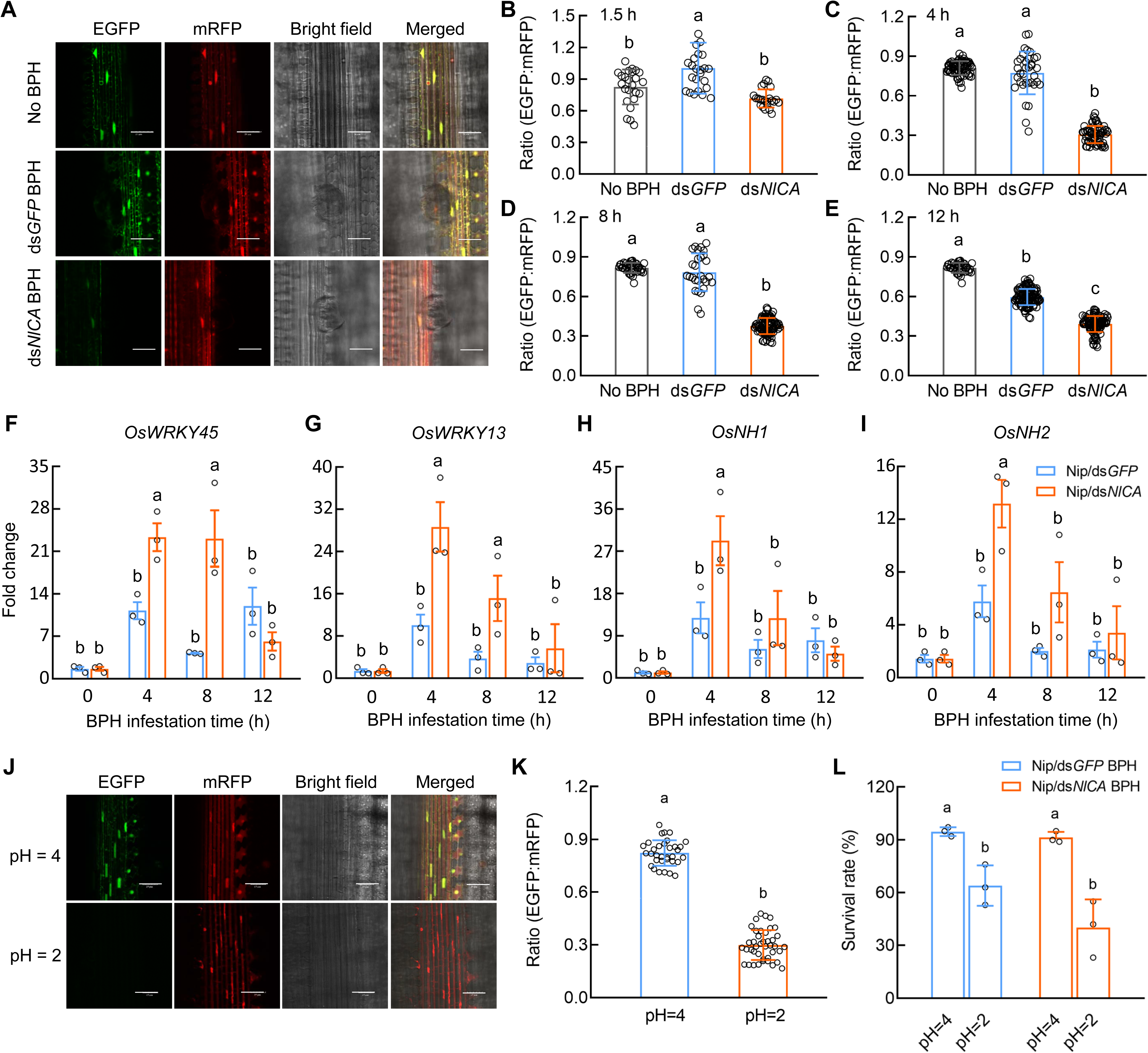
Role of NlCA in maintaining intracellular pH of rice cells. (A) Confocal microscopic images at BPH feeding sites in Nipponbare plants expressing a ratiometric cytoplasmic pH sensor (Cyto-pHusion) 4 h after placing 30 5^th^-instar BPH nymphs on each plant. A 40.0 x objective was used to capture every feeding site on the tiled 1 cm x 1 cm section of rice leaf sheath. Confocal images from plants with no BPH feeding served as the control. Scale bar = 20 μm. (B–E) EGFP:mRFP signal ratios at 1.5 h (B), 4 h (C), 8 h (D) and 12 h (E) of ds*GFP* or ds*NlCA* BPH treatment compared with no BPH control. EGFP was imaged at λ_Ex_ = 500 nm and λ_Em_= 540 nm. mRFP was imaged at λ_Ex_= 570 nm and λ_Em_ = 620 nm. Values are displayed as mean ± SEM (n ≥ 24 circular areas of leaf sheath phloem with a diameter of 100 μm with the feeding site at the center). (F–I) Expression of defense response genes, *OsWRKY45* (F), *OsWRKY13* (G), *OsNH1* (H) and *OsNH2* (I), in Nipponbare plants infested by ds*GFP* or ds*NlCA* BPH. Values are displayed as mean ± SEM of three biological replicates. Each biological replicate represents pooled leaf sheaths from three individual rice plants fed by 20 5^th^ instar BPH nymphs per plant. (J) Confocal microscopic images of Nipponbare plants expressing a ratiometric cytoplasmic pH sensor (Cyto-pHusion) at 12 h after being transferred to Yoshida medium at pH of 2. Confocal images from plants grown in Yoshida medium with pH of 4 served as the control. Scale bar = 15 μm. (K) EGFP:mRFP signal ratios. EGFP was imaged at λ_Ex_ = 500 nm and λ_Em_ = 540 nm. mRFP was imaged at λ_Ex_ = 570 nm and λ_Em_ = 620 nm. Values are displayed as mean ± SEM (n ≥ 33 calculation area per condition). (L) The 7^th^-day survival rates of ds*GFP*- and ds*NlCA*-injected BPH insects feeding on Nipponbare growing in media with different pHs. Values are displayed as mean ± SEM of 3 biological replicates (20 individual insects per each biological replicate). Different letters indicate statistically significant differences analyzed by two-way ANOVA (Tukey test, *P* < 0.05). Experiments were repeated three times with similar trends. See also Figure S2 and Figure S3.

Next, we conducted experiments to determine if intracellular acidification is a highly localized cell-type specific response or a spreading local response. We imaged and calculated the intracellular pH in the mesophyll cells and epidermal cells at the feeding site. We found that intracellular acidification occurred in the mesophyll cells and epidermal cells (Supplemental Figure 2I–2L) in response to feeding by ds*NlCA* BPH, albeit to a lesser degree compared to that in the sieve elements/companion cells. In contrast, rice plants fed by ds*GFP* wild-type BPH responded with an initial slight intracellular acidification at 1.5 h, but then consistently maintained a more alkalized intracellular pH in the mesophyll and epidermal cells compared to non-fed rice plants and ds*NICA* BPH (Supplemental Figure 2I–2L).

Because extracellular pH change is associated with plant responses to biotic stress^25,26^, we speculated that rapid intracellular acidification of ds*NlCA* BPH feeding might induce defense gene expression. To test this possibility, Nipponbare plants infested by ds*GFP* and ds*NlCA* BPH were examined for defense gene expression. Commonly used marker genes (e.g., *OsNH1*, *OsNH2*, *OsWRKY45*, *OsWRKY13*) in the studies of rice defense against BPH were selected^27–30^. We found that expression of those genes was significantly induced at a higher level by ds*NlCA* BPH infestation than by ds*GFP* BPH control at 4 h and 8 h (Figure 3F–I), which is consistent with the intracellular pH change at 4 h and 8 h (Figure 3C–D). These results indicate that intercellular acidification is linked to rice defense activation.

### Ectopic intracellular acidification causes defense gene expression in rice plants

Because plant defense responses are important for limiting BPH survival in rice^28,31–33^, we conducted experiments to determine if intracellular acidification would trigger defense gene expression. Rice is normally grown in Yoshida medium, pH of 4^34^. To test if medium pH changes can modulate defense responses in rice, we first grew rice seedlings in Yoshida medium until they reached the 5-leaf stage and then placed them to fresh Yoshida medium with pH adjusted to 2, 3, 4, 5, 6 and 7, respectively, for 48 hours. Interestingly, drastically increased expression of defense response genes (e.g., *OsNH1*, *OsNH2*, *OsWRKY45*, *OsPBZ1*) was observed in Nipponbare plants at acidic pH of 2 (Supplemental Figure 3A–H). The pHusion sensor plants exposed to external pH of 2 showed intracellular acidification that simulates pH changes observed during BPH feeding (Figure 3J–K). Survival rates of BPH insects were lower in Nipponbare plants growing in media with pH of 2, compared to those growing in media with pH of 4 (Figure 3L). Rice growth medium of pH 2 was used to attain an intracellular pH similar to that during BPH feeding (Figure 3K). These results suggest that cellular acidification is sufficient to induce defense gene expression in rice and to reduce BPH’s ability to survive on rice.

### Rice defense responses are inhibited by NlCA

As NICA stabilizes intracellular pH during WT BPH feeding (Figure 3A–D) and both ds*NlCA* BPH feeding and ectopic intracellular acidification causes activation of defense gene expression (Figure 3F–I; Supplemental Figure 3), we next tested the hypothesis that a major function of NICA-mediated stabilization of intracellular pH may be to prevent over-stimulation of downstream rice defense responses during feeding. Callose deposition in the phloem sieve tubes is a classical defense response that is associated with feeding of piercing-sucking insects^35^. We examined this response using aniline blue to stain callose in the phloem sieve cells. Indeed, we found that fewer and smaller callose deposition was found in the sieve plates of *NlCA*-expressing leaf sheaths compared to those found in the sieve plates of Nipponbare fed by BPH (Figure 4A–H) at 72 h after BPH feeding, demonstrating that *Nl*CA suppresses callose deposition in sieve cell pates. Furthermore, ds*NlCA* BPH induced higher expression of callose biosynthesis genes, such as *OsGSL1*, *OsGSL3*, *OsGLS5* and *OsGns5*, than the ds*GFP* control BPH; however, the induction of these genes was greatly compromised in *NlCA* transgenic plants with or without BPH treatment (Figure 4I–L). We also measured the transcript levels of defense marker genes (e.g., *OsNH1, OsNH2, OsWRKY45* and *OsWRKY13*) in Nipponbare and *NlCA*-expressing plants fed by ds*GFP* and ds*NlCA* BPH. As shown in Figure 4M–P, induced expression of *OsNH1, OsNH2, OsWRKY13* and *OsWRKY45* by ds*NlCA* BPH was suppressed in *NlCA* transgenic plants compared to those in Nipponbare plants. Collectively, these results showed that NICA-mediated intracellular pH homeostasis is linked to downregulation of callose deposition in phloem sieve cells as well as defense gene expression in rice plants.

**Figure 4.**
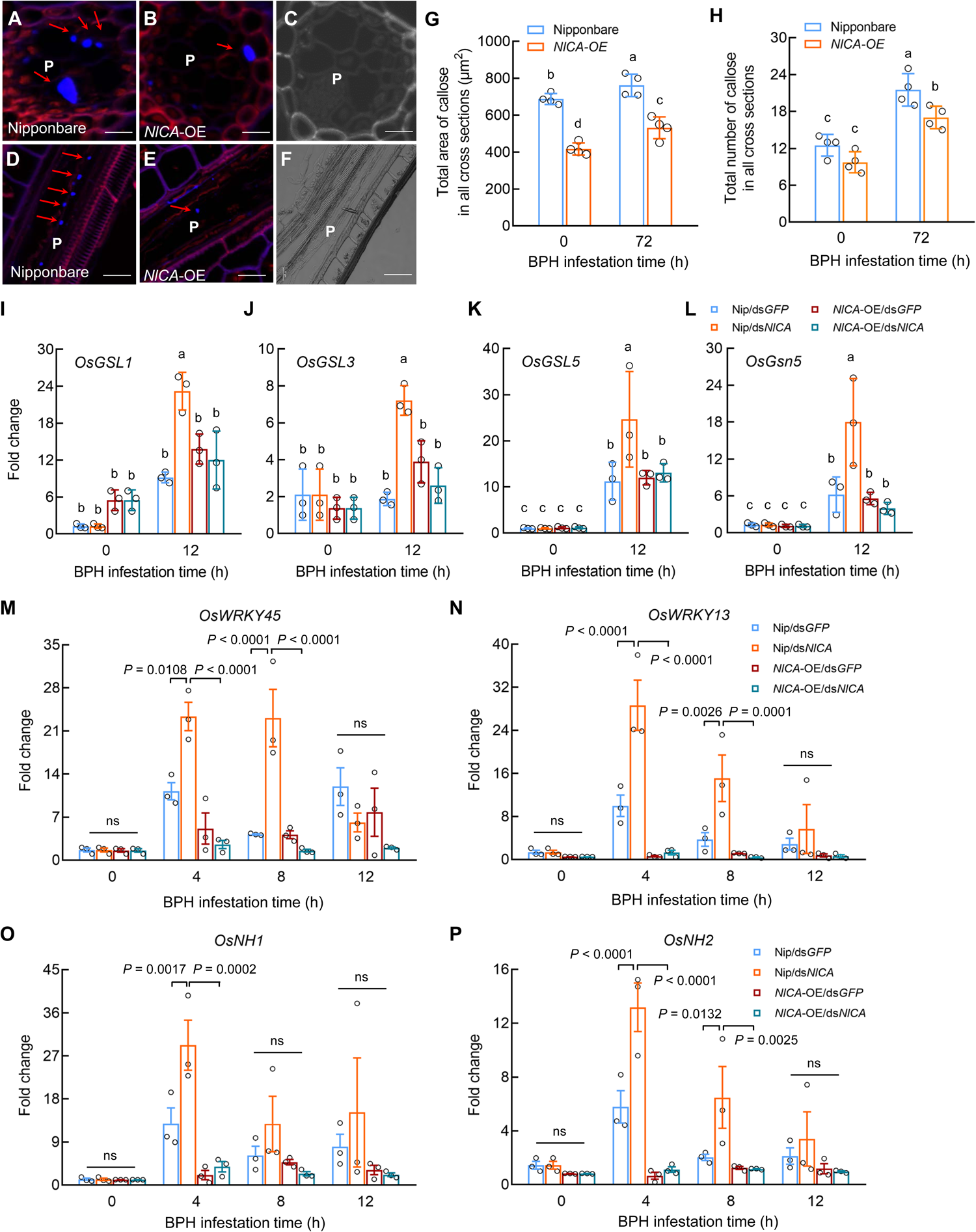
NICA dampens callose deposition and defense gene expression. (A–F) Callose accumulation (bright blue fluorescence indicated by red arrows) on the sieve plates of Nipponbare leaf sheaths (A and D) and *NlCA*-OE leaf sheaths (B and E) plants at 72 h after BPH feeding. P, phloem. The pictures were taken under a Zeiss Microscope. (A–B) Cross-sections. Scale bar = 5 μm. (D–E) Longitudinal sections. Scale bar = 10 μm. (C and F) The bright field views of cross and longitudinal phloem sections, respectively. (G) Total areas of callose deposition in BPH-infested leaf sheaths of Nipponbare and *NlCA*-OE. Each data point represents the total areas of callose deposition found in 300 cross-sections of each experiment. Values are displayed as mean ± SEM of four experiments. (H) Total number of callose deposits in BPH-infested leaf sheaths of Nipponbare and *NlCA*-OE. Each data point represents the total number of callose deposits found in 300 cross-sections of each experiment. Values are displayed as mean ± SEM of four experiments. (I–L) Relative expression levels of the callose synthase gene *OsGSL1* (I), *OsGSL3* (J), *OsGSL5* (K) and *OsGns5* (L) in response to BPH feeding. Values are displayed as mean ± SEM of 3 biological replicates. Each biological replicate represents pooled leaf sheath form three individual rice plants fed by 20 5^th^ instar BPH nymphs per plant. (M–P) Expression of defense marker gene *OsWRKY45* (M), *OsWRKY13* (N), *OsNH1* (O) and *OsNH1* (P) in response to BPH feeding. Values are displayed as mean ± SEM of 3 biological replicates. Each biological replicate represents pooled leaf sheaths from three individual rice plants fed by 20 5^th^ instar BPH nymphs per plant. ns indicates no significant difference between treatments (Two-way ANOVA). Different letters indicate statistically significant differences analyzed by two-way ANOVA (Tukey test, *P* < 0.05). Experiments were repeated three times with similar trends.

## Discussion

In this study, we provided evidence that intracellular acidification is a previously unrecognized plant defense response that occurs during BPH feeding on rice. This finding was facilitated by our attempt to understand the role of NICA in the rice-BPH interaction. We found that the *NlCA* transcript is detected mainly in the salivary glands (Figure 1A) and that the NICA protein is found in rice tissues, including the phloem sap, fed by BPH (Figure 1B). *NlCA*-silenced (ds*NlCA*) insect survived very poorly on at least two independent cultivars of rice plants, Xiushui134^6^ and Nipponbare (Figure 2), whereas *NlCA* expressing rice plants can restore the normal survival of *NlCA*-silenced (ds*NlCA*) insects (Figure 2; Supplemental Figure 1), suggesting that NlCA functions in plant cells. Using the cytoplasm pH sensor, we found that NlCA is required for BPH to maintain a normal plant cytoplasm pH during BPH feeding (Figure 3A–D; Supplemental Figure 2). Pathogen/insect-derived effectors can be powerful molecular probes to discovering novel plant regulators/responses to biotic attacks. Although other piercing/sucking herbivores derived defense-suppressive effectors^36–41^, our discovery of intracellular acidification as a plant defense response and our finding that brown planthopper (BPH) secretes NICA as a counter-defense measure illustrates host cell pH homeostasis as a novel battlefield that has not been revealed in any plant-biotic interactions.

Extracellular alkalinization of culture plant cells has long been recognized as a canonical plant response to microbial elicitors as well as endogenous plant signals^26^. Extracellular alkalinization caused by plant endogenous RAPID ALKALINIZATION FACTOR (RALF) peptides, for example, are perceived by the FERONIA-family receptors^42,43^. There is evidence that this perception causes phosphorylation of PM-localized H(+)-ATPase 2, resulting in the inhibition of proton transport across the PM^44^. Notably, extracellular alkalinization has recently been shown to inhibit or promote growth- or immunity-associated cell surface receptor functions through specific pH-sensitive amino acid sensors^26^. In contrast, defense regulation by intracellular pH changes has so far escaped the discovery by researchers until this study. Our demonstration of a link between intracellular acidification and defense activation has laid a foundation for future discovery of potentially diverse pH-sensitive intracellular regulators of defense responses, which could add a new dimension in the study of plant-biotic interactions.

Because cellular pH alterations could potentially affect multiple biomolecules and, hence, multiple cellular processes, future research should comprehensively define all cellular processes that are affected by intracellular pH acidification. In this study, we found that this pH change is linked to activation of callose deposition at phloem sieve cells and expression of defense response genes, such as *OsNH1, OsNH2, OsPBZ1* and *OsWRKY45* (Figure 3; Supplemental Figure 3). In reverse, NICA-mediated intracellular pH stabilization dampens these defense responses (Figure 4). Callose deposition in phloem sieve cells, in particular, is a classical defense response to a variety of sucking/piecing insects and is thought to limit nutrient flow during insect feeding^35,45–48^. Together, these results suggest that a major effect of NlCA-mediated pH stabilization is to prevent overactivation of defense responses during BPH feeding.

The mechanism by which NICA counters intracellular acidification is likely inherent to its reversible inter-conversion of carbon dioxide and water into carbonic acid, protons and bicarbonate ions. CAs are universally present in all organisms (Supplemental Figure 4); other piecing/sucking insects may use CAs or another mechanism to manipulate host intracellular pH as part of their infestation strategy. Indeed, CA has been reported as a protein component of saliva in rice green leafhopper, *Nephotettix cincticeps*^49^, and aphid *Myzus persicae*^50^. In the case of *M. persicae,* CA-II was shown to increase viral transmission via plant apoplastic acidification-mediated acceleration of intracellular vesicular trafficking^50^. However, because NICA plays a critical role in BPH’s survival on rice plants *per se* (i.e., in the absence of viral infection), as shown in this study, it is more likely that insect-secreted CAs constitute play a primary role in facilitating insect survival by countering intracellular acidification-associated defense activation. In fact, future research should examine if CA-mediated increase in viral transmission may be a secondary consequence of defense suppression, which was not examined in the previous study^50^.

Discovering the role of pH regulation during plant response to biotic and abiotic stresses and characterizing the impact of such pH alterations could be an important area for future research. Because maintaining proper external and internal pH is critical for all forms of life, prokaryotic and eukaryotic organisms, alike, have evolved mechanisms to achieve pH homeostasis. Facing the fluctuating external pH, prokaryotes have evolved diverse mechanisms for sensing external pH. For instance, the bimodal sensing of pH is employed by *Bacillus subtilis* and *Escherichia coli*^51^. Fungi employ a conserved pathway, mediated by Rim101 and PacC, to sense external pH^52^. It has been reported that different subcellular compartments within the plant cell maintain different pH values, presumably as part of carrying out their unique physiological functions^53^. The demonstrated ability of NICA to counter stimulus-dependent pH changes in plant cells could make NICA a useful molecular tool to modulate and broadly understand the effects of pH stabilization on plant signaling and metabolic pathways in different cell types, organelles, and tissues in plants by, for example, targeting NICA expression in specific tissues, cells, or organelles. Additionally, the crucial role of NICA for BPH infestation of rice suggests that NICA is an important target for chemical or trans-kingdom RNAi-based inactivation for the development of novel BPH control strategies in plants.

## Supporting information

Supplemental Information-Version 2

## Acknowledgments

We thank our lab member Richard Hilleary for his comments and suggestions during this work. This study was supported by fundings from the Natural Science Foundation of China (32360082 and 31870259, which supported her research at Michigan State University) and the Yunnan Fundamental Research Projects (202401AS070122 and 202105AC160028) to Y.J.J., Zhejiang University (to support the research of X.Y.Z. at Michigan State University), Howard Hughes Medical Institute to S.Y.H. and the National Key Research and Development Plan in the 14^th^ five-year plan (2021YFD1401100) to C.X.Z.

## Author Contributions

S.Y.H., C.X.Z., Y.J.J. and X.Y.Z. conceptualized and designed this study; Y.J.J., X.Y.Z., S.Q.L., Y.C.X., X.M.L., Y.P.Y., Z.Y.P., L.Z., J.B.L., and H.J.H. performed experiments and analyzed data; C.X.Z. and S.Y.H. analyzed data. Y.J.J., X.Y.Z., C.X.Z. and S.Y.H. wrote the paper with all authors approved the final article.

## STAR METHODS

Detailed methods are provided in the online version of this paper and include the following:

- KEY RESOURCES TABLE
- RESOURCE AVAILABILITY

- Lead contact
- Materials availability
- EXPERIMENTAL MODEL AND SUBJECT DETAILS

- Plant materials and insects
- METHOD DETAILS

- Plant materials and insects
- Transgenic rice
- *In situ* mRNA hybridization analysis
- Rice sheath sample preparation for LC-MS
- RNA interference (RNAi)
- BPH survival test on artificial diet
- BPH survival test on rice
- pH assay
- RNA isolation and quantitative real-time PCR (qRT-PCR).
- Callose deposition in rice sheath
- QUANTIFICATION AND STATISTICAL ANALYSIS

## SUPPLEMENTAL INFORMATION

**Supplemental Figure 1.** Generation of *NlCA-OE* and mutant *NlCA-OE* plants, and survival rates of BPH, related to Figure 2.

**Supplemental Figure 2.** Quantification of fluorescent signals in leaf sheaths of Nipponbare plants expressing the cyto-pHusion ratiometric pH sensor in response to BPH feeding, related to Figure 3.

**Supplemental Figure 3.** Defense response genes are modulated by ectopic pH manipulation in rice, related to Figure 3.

**Supplemental Figure 4.** Phylogenic analysis of carbonic anhydrase proteins from representative bacteria, insect, mammal, and plant species, related to Discussion section.

**Supplemental Table 1.** List of detected BPH-secreted salivary proteins, related to Figure 1.

