## Supplemental Information-Version 2 for "Rapid intracellular acidification is a novel plant defense response countered by the brown planthopper"

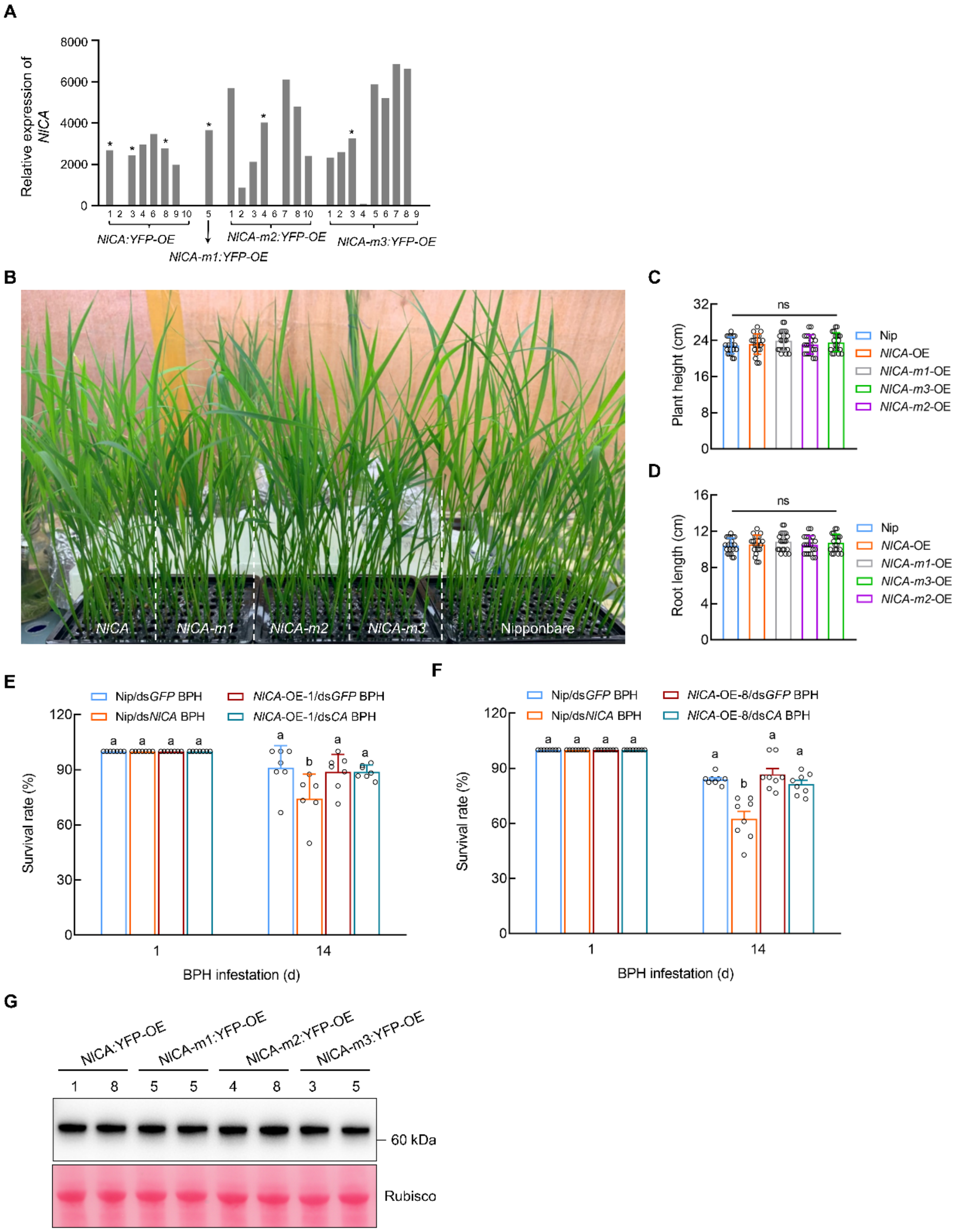
A
Relative expression of NlCA
8000
6000
4000
2000
0
NlCA:YFP-OE
NlCA-m1:YFP-OE
NlCA-m2:YFP-OE
NlCA-m3:YFP-OE
B
NlCA
NlCA-m1
NlCA-m2
NlCA-m3
Nipponbare
C
Plant height (cm)
ns
Nip
NlCA-OE
NlCA-m1-OE
NlCA-m3-OE
NlCA-m2-OE
D
Root length (cm)
ns
E
Nip/dsGFP BPH
Nip/dsNlCA BPH
NlCA-OE-1/dsGFP BPH
NlCA-OE-1/dsCA BPH
Survival rate (%)
BPH infestation (d)
F
Nip/dsGFP BPH
Nip/dsNlCA BPH
NlCA-OE-8/dsGFP BPH
NlCA-OE-8/dsCA BPH
Survival rate (%)
BPH infestation (d)
G
NlCA:YFP-OE
NlCA-m1:YFP-OE
NlCA-m2:YFP-OE
NlCA-m3:YFP-OE
1 8 5 5 4 8 3 5
60 kDa
Rubisco

**H**

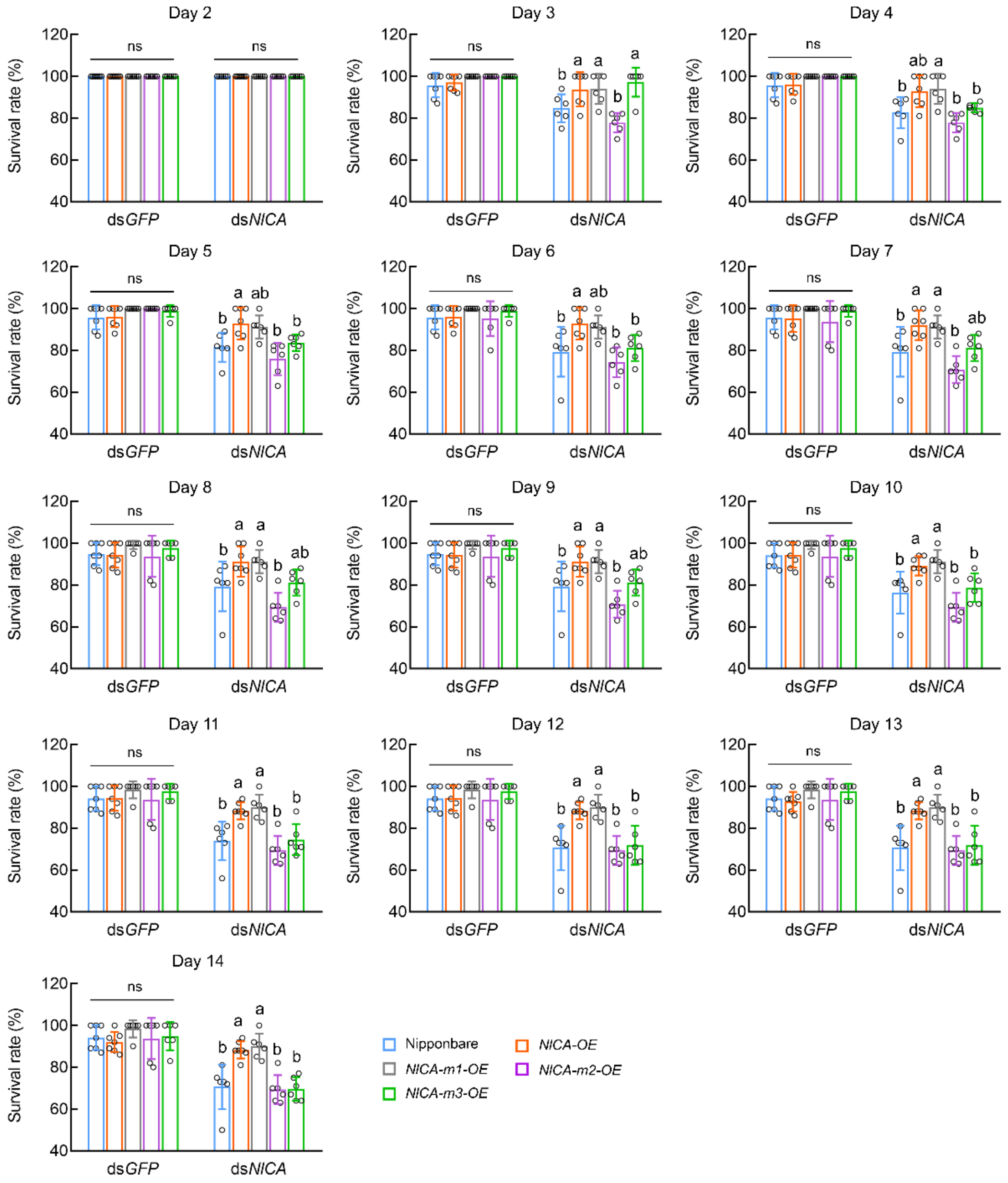


**Supplemental Figure 1**. Generation of *NlCA-OE* and mutant *NlCA-OE* plants and survival rates of BPH, related to Fig. 2. (A) qRT-PCR analysis of the *NlCA* transcript levels in *NlCA-OE, NlCA-m1-OE, NlCA-m2-OE NlCA-m3-OE* plants. Plant lines labeled with an asterisk were selected for further BPH feeding assay. (B) 21-day-old of Nipponbare, *NlCA-OE, NlCA-m1-OE, NlCA-m2-OE NlCA-m3-OE* plants grown in the modified Yoshida medium. *NlCA-OE, NlCA-m1-OE, NlCA-m2-OE NlCA-m3-OE* plants displayed normal appearance compared with Nipponbare. (C and D) Plant height (C) and root length (D), respectively, in (B). Values are displayed as mean ± SD (n = 20 plants). ns indicate no significant statistically differences analyzed by one-way ANOVA (Tukey test). (E and F) The survival rates of ds*GFP* and ds*NlCA* BPH insects on Nipponbare and *NlCA* transgenic plant line 3 (E) and line 8 (F) after 1-day and 14-days post-infestation. Values are displayed as mean ± SEM (n = 8 biological replicates; 20

individual insects per each biological replicate). (G) Protein samples were extracted from one-week-old seedlings of *NlCA*-OE (lines 1 and 8) and *NlCA*-mutant-OE (line 5 of NlCA-m1, lines 4 and 8 of NlCA-m2 and lines 3 and 5 of NlCA-m3) plants. Abundance of fusion proteins were detected with anti-GFP (Abmart, 20004). Ponceau S staining of Rubisco confirmed equal loading. (H) The survival rates of ds*GFP* and ds*NlCA* BPH insects on Nipponbare and *NlCA* transgenic plants. Values are represented as mean ± SEM (n > 6 biological replicates; 20 individual insects per each biological replicate). Different letters indicate statistically significant differences analyzed by two-way ANOVA (Tukey test, $P < 0.05$). Experiments were repeated three times with similar trends.

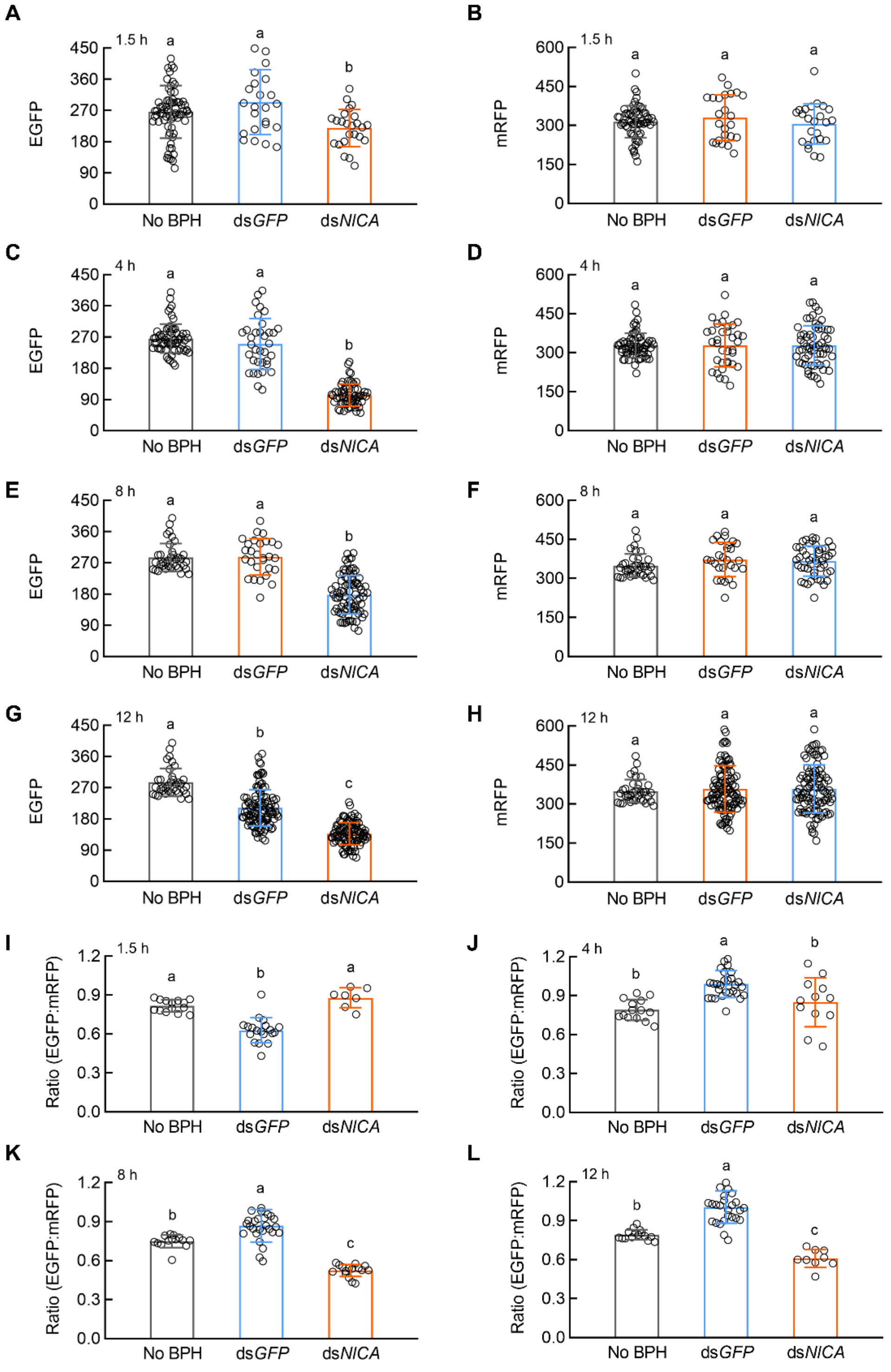


**Supplemental Figure 2**. Quantification of fluorescent signals in leaf sheaths of Nipponbare plants expressing the cyto-pHusion ratiometric pH sensor in response to BPH feeding, related to Fig. 3. (A, C, E, G) eGFP fluorescence intensity at 1.5, 4, 8 and 12 h after BPH feeding. (B, D, F, H) mRFP fluorescence intensity at 1.5, 4, 8 and 12 h after BPH feeding. Values are displayed as mean ± SEM (n > 14 feeding sites for each genotype). (I–L) Intracellular acidification of the mesophyll cells and epidermal cells at BPH feeding sites. EGFP:mRFP signal ratios at 1.5 h (I), 4 h (J), 8 h (K) and 12 h (L) of ds*GFP* or ds*NlCA* BPH feeding compared with no BPH control. EGFP was imaged at $\lambda_{Ex}$ = 500 nm and $\lambda_{Em}$ = 540 nm. mRFP was imaged at $\lambda_{Ex}$ = 570 nm and $\lambda_{Em}$ = 620 nm. Values are displayed as mean ± SEM (n ≥ 16 circular areas of leaf sheath, each circular area had a diameter of 100 μm with the feeding site at the center). Image processing and analysis was performed by Fiji. Different letters indicate statistically significant differences analyzed by one-way ANOVA (Tukey test, $P < 0.05$).

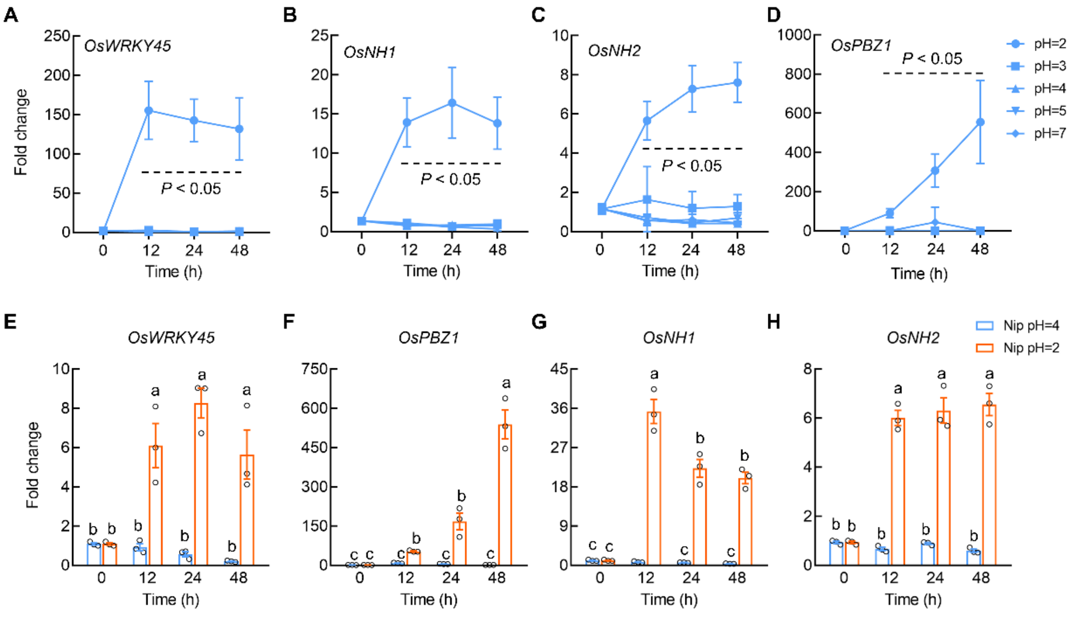


**Supplemental Figure 3**. Defense response genes are modulated by ectopic pH manipulation in rice, related to Fig. 3. (A-D) Expression levels of defense response genes, *OsWRKY45* (A), *OsNH1* (B), *OsNH2* (C) and *OsPBZ1* (D), are induced after acidification of Yoshida medium in which WT Nipponbare plants were grown. Rice plants were first grown in Yoshida media with pH of 4 to 5-leaf-stage and were then placed into fresh Yoshida medium with different pH for 48 h. RNA samples collected from the stem tissues for RT-qPCR in the indicated time. (E-H) Expression levels of defense response genes, *OsWRKY45* (E), *OsPBZ1* (F), *OsNH1* (G) and *OsNH2* (H), are induced after acidification of Yoshida medium in which WT Nipponbare plants were grown. Rice plants were first grown in Yoshida media with pH of 4 to 5-leaf-stage and were then placed into fresh Yoshida medium with pH of 4 or pH of 2 for 48 h. RNA samples collected from the stem tissues for RT-qPCR in the indicated time. Values represent mean ± SEM (Two-way ANOVA, n = 4). Different letters indicate statistically significant differences analyzed by two-way ANOVA (Tukey test, $P < 0.05$). Experiments were repeated three times with similar trends.

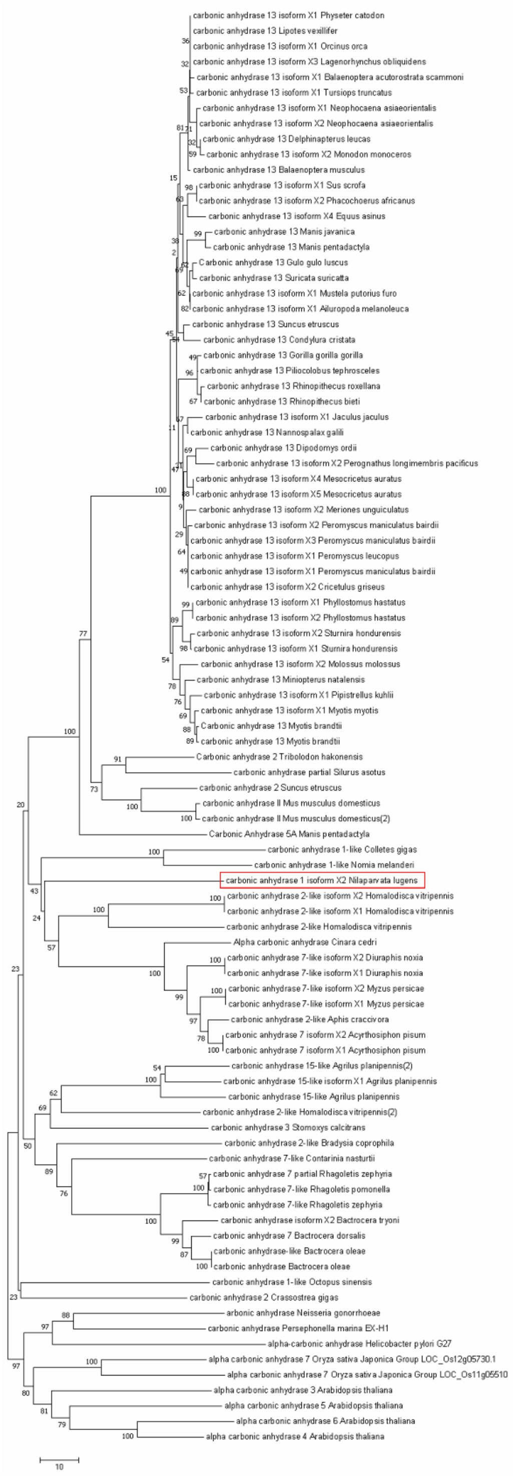


**Supplemental Figure 4. Phylogenic analysis of carbonic anhydrase proteins from representative bacteria, insect, mammal, and plant species, related to Discussion section.** The protein sequences were obtained from NCBI (www.ncbi.nlm.nih.gov), TAIR (http://www.arabidopsis.org/) and Rice Genome Annotation Project (http://rice.uga.edu/index.shtml). Red rectangle denotes NlCA. Sequence alignment was performed using Clustal W, and the phylogenetic tree was generated by MEGA 7.0 using the Neighbor–Joining method. Bootstrap values from 1,000 replicates were used to assess the robustness of the tree. The scale indicates the average number of substitutions per site.
